# A major life-history locus underlies genotype-by-environment variation in growth across water temperatures

**DOI:** 10.64898/2026.05.06.723333

**Authors:** Ana Sofia Lindeza, Corinne Suvanto, Gautier Magne, João Lopes, Amine Ejite, Morgane Frapin, Anti Kause, Craig Primmer

## Abstract

Rapid environmental change is exposing organisms to conditions that do not match those under which they evolved, making it increasingly important to understand how genetic variation shapes phenotypic responses to environmental stress. Since most phenotypic traits arise from interactions between genetic variation and the environments experienced throughout an organism’s lifetime, understanding the genetic architecture of these interactions is central to predicting how populations will respond to novel environments. While genotype-by-environment interactions (G×E) are well studied in quantitative genetics, identifying specific loci that contribute to environmentally dependent trait expression remains rare. Salmonids already exhibit a wide portfolio of plastic life-history strategies, reflecting adaptation to highly heterogeneous environments, yet it remains unclear whether known major-effect loci involved in life-history regulation also contribute to variation in plastic responses to environmental change. One such major-effect locus is the transcription factor *six6*, which has been repeatedly associated with variation in age at maturity across multiple populations of rainbow trout (*Oncorhynchus mykiss*). Since maturation timing is closely linked to growth trajectories and patterns of energetic allocation during early development, allelic variation at this locus may also influence growth responses to warming conditions. Here, we test this hypothesis using a common-garden experiment in which 6 months old juvenile rainbow trout were reared under current and warming (+2°C) temperature regimes. By quantifying genotype-specific reaction norms across environments, we show that *six6* genotype contributes to environmentally dependent variation in growth and body composition, with individuals heterozygous for the *six6* locus showing a distinct and steeper response to warming relative to homozygotes. These findings provide evidence that a major life-history gene shapes plastic responses to thermal stress in juvenile rainbow trout, with novel implications for how standing genetic variation at in large-effect loci may influence population-level responses to climate warming.

## 1. Introduction

Anthropogenic climate change and rapidly fluctuating environments are pushing organisms to respond and adapt to conditions of no historical precedent (Hoffmann & Sgrò, 2021). The recorded, and planned to increase, changes in temperature pose serious risks to populations persistence, growth and reproductive fitness (Pörtner & Farrel, 2008). Although continents are warming faster than oceans, the strongest temperature increases are projected for the northernmost regions of the world, where terrestrial and water systems are experiencing a concerningly rapid warming (IPCC, 2024). Rising temperatures imply that many species will experience thermal conditions that no longer match the environment under which they evolved and were selected, leading to cue mismatch and, in extreme cases, to population decline or local extinction (Parmesan, 2006; Reed et al., 2010; Chevin et al., 2010; Urban, 2015). The consequences of the rising temperatures for fish populations are already evident with documented effects on migration timing, feeding behavior, metabolic performance, and other fitness-related traits (Volkoff & Rønnestad, 2020; Jonsson, 2023). Salmonids are known to be particularly sensitive to environmental change and, as ectothermic animals with complex life cycles spanning multiple environments, have their performance and survival tightly linked to temperature conditions (Jonsson, 2023; Ahi et al., 2025). Like many organisms facing rapid environmental change, salmonids can respond to these changes by either shifting their distribution to find better environmental conditions or by adjusting their phenotype to the new environment. Because contemporary climate change is occurring on timescales far shorter than those required for adaptive evolution through new mutations, short-term responses are expected to occur primarily through phenotypic plasticity (Chevin et al., 2010; Reed et al., 2010). Phenotypic plasticity allows a single genotype to produce different phenotypes depending on environmental conditions, thereby modulating how populations respond to environmental variation and even buffering them against rapid environmental change (Chevin et al., 2010; Merilä & Hendry, 2014). Variation in plastic responses among individuals, or genotypes, is commonly described as genotype-by-environment (G×E) interaction and simply defined as the mean phenotypic changes of a given genotype in different environments, often visualized as differences in reaction norms (Oomen & Hutchings, 2022).

Most studies focus mainly on plastic responses during the marine phase of salmonid life cycles, often related to migration behavior, marine survival or maturation timing (Jonsson et al., 2003; Friedland et al., 2005; Russel et al., 2012; Barson et al. 2015; Vehanen, et al., 2023), but rarely prioritize the environmental conditions experienced during early freshwater development. Yet early life-history traits are known to demonstrate remarkable flexibility in the face of environmental change responding through both genetic and non-genetic mechanisms, but the magnitude and nature of these responses can vary substantially across salmonid species (Jonsson & Jonsson, 2009; Jonsson and Jonsson, 2014; Scholl et at, 2026).

Furthermore, it is well accepted that ecological changes in one life stage can have extensive consequences for later life stages, because the various life-stage transitions are finely tuned to conditions in very different environment (Tuljapurkar et al., 2009; Jonsson and Jonsson, 2014; Le Coeur et al., 2022). Since freshwater ecosystems are expected to undergo stronger and more rapid thermal changes than marine environments, due to their smaller area, the early life stages of salmonids are likely to experience particularly strong thermal variation, leading to downstream consequences for later life-history transitions and creating conditions under which phenotypic plasticity and GxE might play a key role in shaping developmental and life history trajectories (Crozier & Hutchings, 2014; Oomen & Hutchings, 2022).

For salmonids, such as rainbow trout (*Oncorhynchus mykiss*), studies have shown that early growth trajectories are particularly sensitive to environmental temperature, since thermal conditions experienced during development heavily influence muscle cellularity, development and overall growth (Wilkes et al., 2001; Dubey et al., 2023). Since early growth patterns can generate differences in performance and timing of life-history transitions, early life stages are particularly relevant for understanding plastic and genotype-by-environment responses. Although G×E interactions have been widely documented for multiple traits in adult rainbow trout (Sae-Lim et al., 2013; 2015; Janhunen et al., 2016; Gallardo-Hidalgo et al., 2021), most studies have characterized plasticity at the level of populations or quantitative genetic variance components, leaving the genetic architecture of underlying plastic responses to environmental variation largely unresolved. As most complex traits are influenced by many loci with small effects, the real difficult lies in identifying genes that directly contribute to G×E interactions, where the effect of a particular genotype changes across environments, and moving from describing polygenic genotype-by-environment interaction typically captured as population or individual-level reaction norm (MacKay et al., 2009). However, in the case of rainbow trout, some major-effect loci known to influence key life-history traits have been identified for several complex traits, providing a rare opportunity to investigate locus-specific GxE interactions. One such locus is the transcription factor *six6*, a partially conserved biallelic gene with alleles associated with early (E) and late (L) maturation and repeatedly linked to age at maturity across multiple populations of rainbow trout (Sinclair-Waters et al., 2020; Waters et al., 2021; Willis et al., 2020). Genetic variation at *six6* has also been associated to other key life-history traits across salmonids, including spawning ecotype differentiation or migration timing (Cauwelier et al., 2018; Sinclair-Waters et al., 2020; 2022; Moustakas-Verho et al., 2020, Willis et al., 2020 Waters et al., 2021), and with new emerging evidence of its influence in early-life survival dependent on nutrients availability (Aykanat et al., 2024). Whether major-effect loci associated with life-history timing also contribute to variation in plastic responses to environmental change therefore remains an important open question (Doctor et al., 2014).

Using a controlled common-garden experiment with four full-sib families reared under control and warming (+2°C) conditions, we tested whether variation at the six6 locus is associated with genotype-by-environment interactions in early-life growth in juvenile rainbow trout. Fish were reared for six months under two temperature regimes representing control conditions and a warming scenario (+2°C), with measurements taken at three time points (winter, spring and early summer) to quantify growth trajectories, including body length, weight, and body condition. We found a GxE for body weight, already detectable in spring and getting more pronounced towards early summer, with *six6*\*EE individuals showing a much steeper reaction norm between the two environments and strong temperature-dependent differences in growth. Heterozygous, *six6*\*EL, individuals showed the weakest response to temperature, with reduced growth under warming and the flattest reaction norm across treatments. To our knowledge, these results provide the first evidence of locus-specific GxE for early life-history traits in rainbow trout and offer a possible link between genetic variation and phenotypic plasticity, highlighting how responses to warming environments may differ among genotypes within populations.

## 2. Material and Methods

### 2.1. Fish crossing and rearing

Gametes from a set of parental fish, genotyped for a panel of key life-history loci, were obtained from a hatchery in Enonkoski operated by the Natural Resources Institute Finland (LUKE). The broodstock, from which the parental generation subset derives, is part of the Finnish national rainbow trout breeding program and originates from a line established in the 1980s from multiple North American populations, including both freshwater and anadromous lineages (Siitonen, 1986). All selected parental individuals were heterozygous at the *six6* locus, ensuring that offspring segregated all three genotypes (*six6\**EE, *six6\**EL, *six6*\*LL) in Mendelian proportions. In mid-May 2024, eggs and milt were stripped from selected individuals and transported to the University of Helsinki, where artificial fertilizations were conducted the following day. This produced a total of 12 full-sib families, of which four were selected for the present experiment. Eggs and alevins were incubated in vertical trays, under constant dark conditions and water flow. Water temperature was initially set for 7 °C and gradually increased to 10°C in the period of one week, by an increment of 0.5 °C per day, after which it remained constant for the entire incubation period. In July 2024, approximately two months post-hatching, juveniles were transferred to the Lammi Biological Station and reared under standard conditions for three months. Each family was maintained in a separate tank (45 × 94 cm) until the start of the experiment, supplied with flow-through water from Lake Pääjärvi (see Åsheim et al., 2021 for details). Fish were fed a commercial diet (Alltech Fennoaqua) *ad libitum*, with feeding rates adjusted bi-monthly based on tank biomass estimates.

### 2.2. Family genotyping

In October 2024, approximately 3700 individuals from the four selected full-sib families were anesthetized using buffered tricaine methanesulphonate (MS-222; Merck) and tagged with a 8mm passive integrated transponder (PIT) tags for individual identification (Biomark). A small caudal fin clip was collected from every individual for DNA extraction and subsequent genotyping using the Kompetitive allele-specific PCR assay, or KASP (He et al., 2014). Fin clips were placed directly into 96-well plates containing 25 µL of Lucigen QuickExtract DNA Extraction Solution 1.0 and kept on ice or at -20°C until extractions were completed, according to manufacturer’s instructions. The targeted SNPs, in all the individuals, were *six6*, on chromosome 25, and *sdY*, a master sex-determining gene. For the *six6* locus, assays were designed by LGC Biosearch Technologies using two allele-specific forward primers (CTACACGCTTGTCTTGAACTGATTCGT and ACGCTTGTCTTGAACTGATTCG) and and one common reverse primer (GCATAAAATAACCCTGTGATAAACTAACAT). The SDY locus was genotyped using an amplification/non-amplification assay targeting the male-specific region (Sinclair-Waters et al., 2022). Each reaction consisted of 2.5 µL of genomic DNA, 2.5 µL of KASP 2× Master Mix, and 0.07 µL of KASP Assay Mix containing locus-specific primers. PCR reactions were performed on a Bio-Rad CFX Maestro instrument following the protocol described by Sinclair-Waters et al. (2020). Genotypes were automatically called based on allele-specific fluorescence (FAM and HEX).

### 2.3. Experimental design and data collection

By early November 2024, a subset of individuals was selected from each of the four full-sib families based on sex and genotype at the *six6* locus. Approximately 200 individuals were selected from each of the families, with the three *six6* genotypes (EE, EL, LL) represented in comparable proportions across thermal treatments. Families were evenly distributed between treatments, minimizing potential confounding effects between family, genotype and treatment. Individuals were identified using their PIT tag codes, anesthetized with buffered MS-222, and measured for weight (g) and fork length (mm; hereafter “length”) prior to tank allocation. Each treatment was replicated across the two tanks, to account for potential tank effects and the allocation of fish across tanks was done while maintaining a similar representation of *six6* genotypes and sex in each tank. Final sample sizes were 205 fish in of Control tank 1 (CT1), 211 in Control tank 2 (CT2), 211 fish in Warm tank 1 (WT1) and 205 fish in Warm tank 2 (WT2), resulting in a total of 416 individuals per treatment and a total of 832 fish used in the entire experiment. Fish were randomly assigned to one of four tanks distributed across two thermal treatments, a control treatment and a warming treatment, in which water temperature was maintained on average 2°C higher. Water temperatures in the control treatment followed the natural seasonal dynamics of Lake Pääjärvi (Figure 1), with temperatures ranging from a minimum of 1.60 to a maximum of 5.82°C and ranging from 1.54 to 8.16°C in the warming treatment. Temperature loggers in each tank recorded water temperature at 30-minute intervals, from which daily means were calculated.

**Figure 1:**
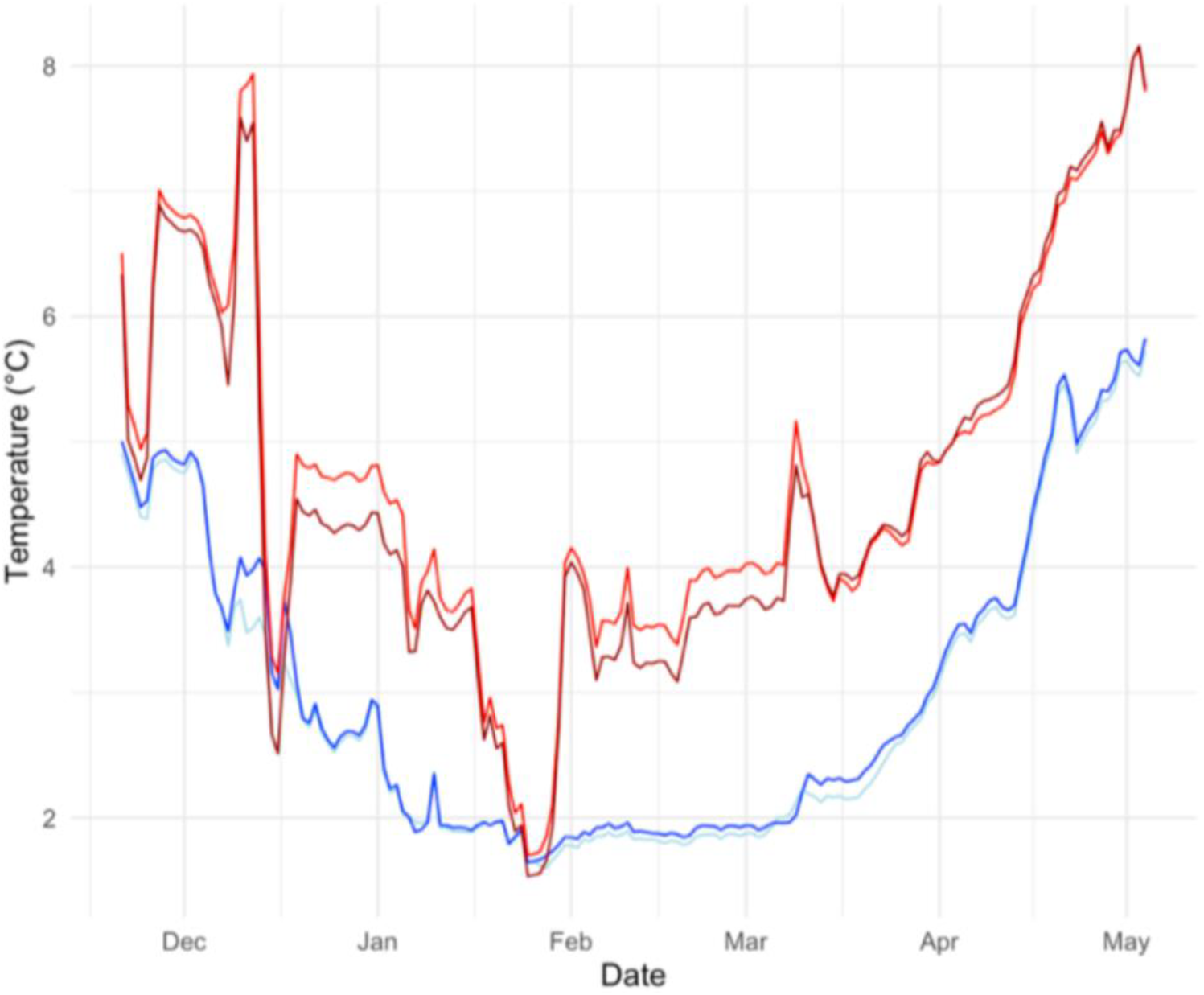
Temperature profiles of experimental treatments. Daily mean water temperatures in the control (blue) and warming (red) treatments from mid-November 2024 to May 2025, reflecting the natural seasonal temperature dynamics of Lake Pääjärvi. In the warming treatment, lake water was heated to maintain an average temperature difference of approximately 2 °C relative to the control. Temperature loggers in each tank recorded measurements at 30-minute intervals, from which daily means were calculated. Temporary decreases in the warming treatment reflect short-term heating system interruptions.

A second measurement was performed in March 2025 to capture the effect of early spring conditions, characterized by increasing day length and rising water temperatures, on the divergence of growth trajectories following the winter period. Tanks were processed sequentially by anesthetizing fish with buffered MS-222, recording their weight and length, and subsequently returning them to their original tanks.

A third and final measurement was carried out in May 2025, characterized by early summer conditions with longer photoperiod and warmer temperatures. Fish were euthanized using buffered MS-222, and weight and length were once again recorded. Afterwards, all the individuals were scanned, using dual-energy X-ray absorptiometry (DXA; iNsiGHT VET DXA, OsteoSys Co., Ltd., Korea), which provides high-resolution estimates of tissue composition by partitioning each scan into fat mass (g), lean (g) mass, and bone mineral content. Derived traits included total fat mass (g), fat percentage, lean body mass, bone area, and bone mineral content (BMC). Body condition was calculated, at each of the three time points, using weight and length measurement using Fulton’s condition factor (K), as 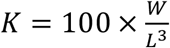 where *W* is body mass (g) and *L* is fork length (mm).

### 2.4 Statistical analysis

All statistical analyses were conducted in R (version 2026.01.1). To test for differences in growth between treatments, we fitted a linear mixed-effects model for body weight, length and condition, at each sampling time point (November, March, and May) using the ‘lme4’ package. For each trait, models included treatment, sex, and their interaction as fixed effects, and family and tank as random intercepts. Family was included as a random effect in all the models, since comparisons based on AIC indicated a better fit compared to treating family as a fixed effect. The model structure was:

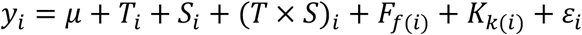

where *y*_*i*_ represents body weight, length and condition of individual *i* at any of the measured time points. To test if the effect of temperature differed between sexes we added an interaction term. Models were fitted separately for each sampling period and trait. Statistical significance of fixed effects, including interaction terms, was evaluated using analysis of variance on fitted models. Because no significant *T* × *S* interaction was detected in the initial models, sex was not included in follow-up analyses, as it did not change the response to temperature.

To test whether growth responses to temperature differed among *six6* genotypes, we fitted the same linear mixed-effects models for body weight, length and condition at each sampling time point using the ‘lme4’ package. Models included treatment, *six6* genotype, and their interaction as fixed effects, and family and tank as random intercepts. The model structure was:

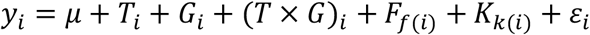

where *G*_*i*_ is the *six6* genotype, treated as a three-level categorical variable (EE, EL, LL). The interaction term between treatment and six6 genotype was used to test for genotype-by-environment (G×E) interactions. Once again, models were fitted separately for each sampling period to assess how G×E effects developed over time. To quantify genotype-specific responses to temperature, we estimated marginal means using the ‘emmeans’ package. These were used to construct reaction norms for each genotype across environments and to estimate pairwise contrasts of treatment effects within genotypes. We then fitted additional models including body length as a covariate, to test whether genotype-dependent differences in body weight were independent of overall body size.

To assess whether genotype-by-environment effects on growth were associated with differences in body composition, we analyzed the DXA derived traits, measured at the final sampling point, including total fat mass, fat percentage, lean mass, bone mineral density (BMD) and bone mineral content (BMC). For each trait, we fitted linear mixed-effects models as:

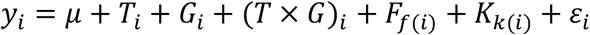

Where *y*_*i*_ is one of the DXA focal traits, *T*_*i*_ is treatment and *G*_*i*_ is the *six6* genotype. Family, *F*_*f*(*i*)_, and tank, *K*_*k*(*i*)_, were included once again as random intercepts. To evaluate whether genotype-by-environment effects on body weight were driven by differences in overall body size, we fitted extended models including body length as a covariate:

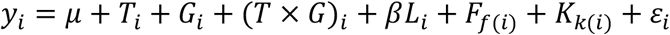

Here *L*_*i*_represents body length. Similarly, to assess whether differences in body composition were independent of body size, we included log-transformed body weight as a covariate in models of DXA-derived traits:

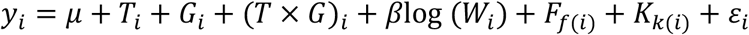

where *W*_*i*_ is body weight. These models allowed us to distinguish whether observed effects were driven by overall size scaling or by differences in body composition independent of size.

To summarise multivariate patterns in body composition independent of size, we performed a principal component analysis (PCA), on size-corrected traits. Each trait was log-transformed and regressed against the log-transformed body length:

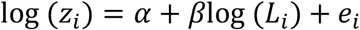

where *z*_*i*_ is the focal trait and *e*_*i*_ are residuals, for size-corrected values. The PCA was then performed on the centred and scaled residuals using the *‘prcomp’* function. Broad-sense heritability (*H*^*2*^) of body weight was estimated separately for each thermal environment using a linear mixed model with fixed effects of sex and *six6* genotype, and random effects of family and tank. *H*^*2*^ was calculated as the proportion of total phenotypic variance attributable to family.

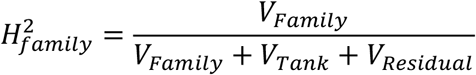

Where *V*_*Family*_represents variance attributable to family, *V*_*Tank*_ represents variance among tanks, and *V*_*Residual*_ is the residual variance. Because pedigree-based additive genetic effects were not explicitly modelled, these estimates represent family-level broad-sense heritability (*H*^*2*^_family_) rather than narrow-sense heritability. Parametric bootstrap confidence intervals (999 iterations) were obtained using the ‘*bootMer’* function, conditioning on observed random effects, which is appropriate given the number of families. The significance of the family variance component was assessed using a one-sided likelihood ratio test comparing models with and without the family random effect. Whether H^2^ differed between thermal environments was assessed by computing the bootstrap distribution of the difference in H^2^ estimates.

## 3. Results

### 3.1. Effect of increased temperature on body weight, length and sex

As expected, body weight (p = 0.78) and length (p = 0.90) did not differ between treatments at the initial sampling point, in November (Fig. 2A and D). However, treatment effects became evident over time, with fish kept in warm condition becoming both heavier and longer, than those in the control treatment, already by March (p < 0.001, Fig.2B and E) and May (p = 0.002, Fig.2C and F). By the end of the experiment, fish in the warm treatment were on average 24g heavier and 3cm longer than those in the control group. Importantly, there was no strong evidence for a *treatment × sex* interaction throughout the experiment, indicating that the effect of temperature on growth did not strongly differ between sexes. Although a weak effect of sex on body weight was detected at later timepoints, in March (p = 0.050) and May (p = 0.044), differences between males and females were small relative to the overall variation. Further, there was no effect of sex on body length at any sampled timepoint (all p > 0.19). Body condition showed no consistent differences between treatments, or sexes, across sampling points and no interaction effects were detected at any time point (all p > 0.05; Table S1).

**Figure 2:**
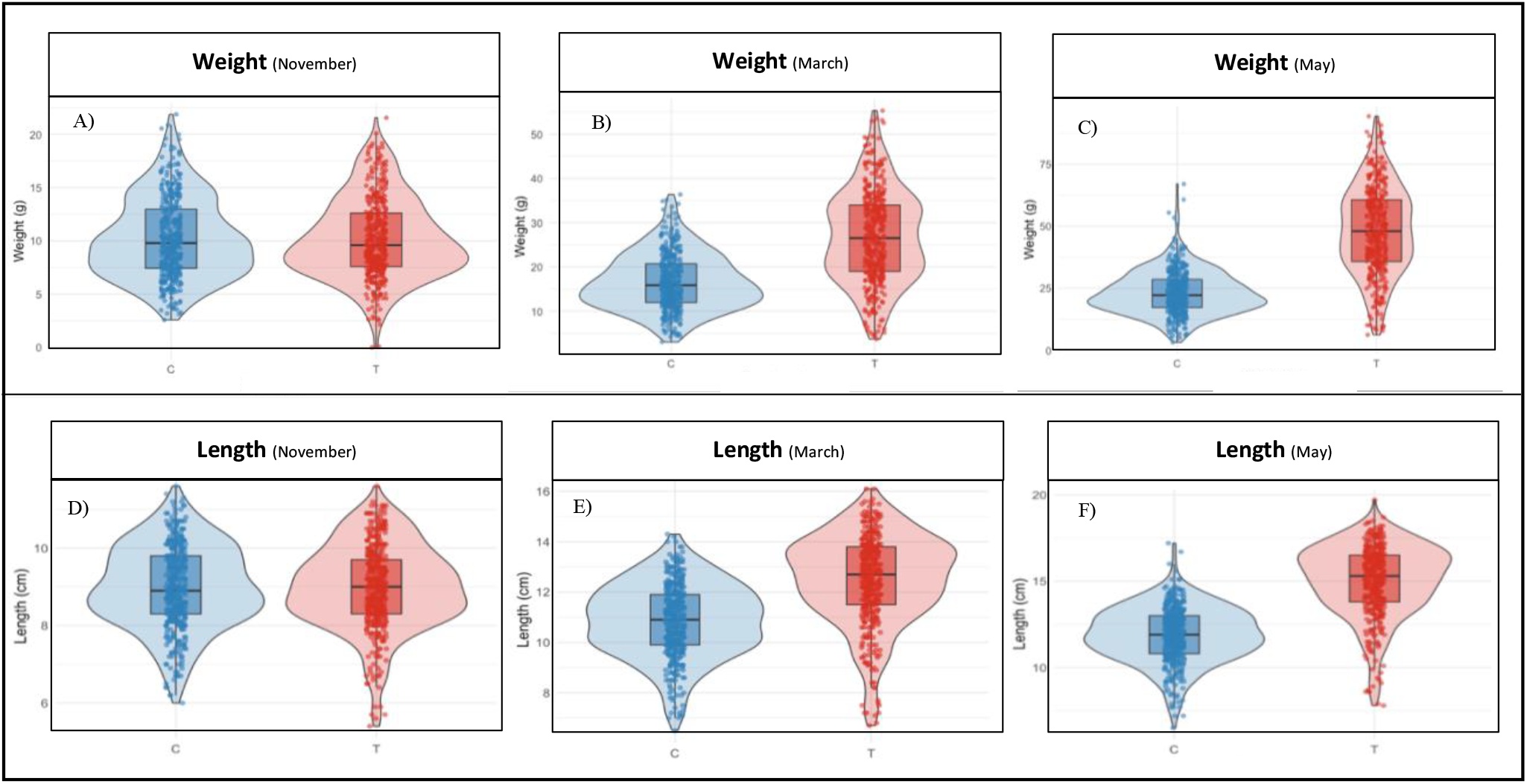
Temperature effects on juvenile rainbow trout growth, from November to May. Body weight (top row) and body length (bottom row) of individuals reared under control temperature (in blue) and a warm environment, increased by 2°C (in red). The experiment had the duration of six months, and body traits were measured in November (left), March (middle) and May (right). Distributions are shown using violin plots with overlaid boxplots and individual data points.No differences between treatments were observed at the initial sampling point (November). By March and May, fish reared under warm conditions exhibited substantially higher body weight and length compared to controls.

**Figure 3:**
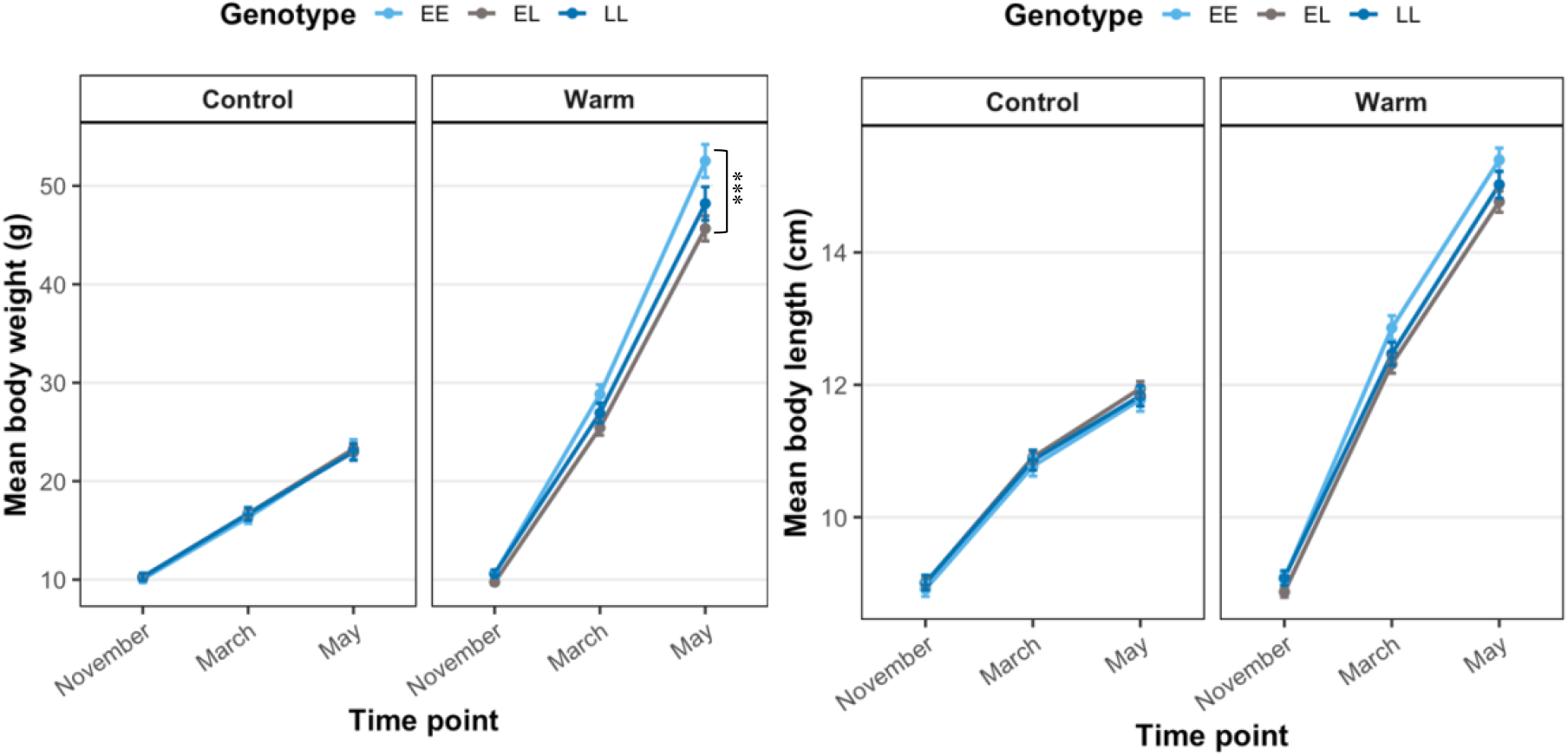
Growth trajectories across time under control vs. warm environments. Mean body weight (A) and mean body length (B) of fish reared under control and warm conditions, across three sampling time point (November, March and May). Lines represent each of the *six6* genotypes (EE, EL and LL) and connect their mean values. Across both traits, fish reared under warm conditions exhibited substantially greater growth compared to those in the control treatment, with differences emerging from March and increasing by May. Genotype-specific trajectories were largely parallel during early development, indicating similar initial growth patterns across six6 genotypes. While this figure illustrates overall growth dynamics, it does not explicitly capture differences in genotype-specific responses to temperature, which are examined using reaction norms in Figure 4.

### 3.2. Temperature-dependent growth differences among *six6* genotypes

To test whether growth responses to temperature differed among *six6* genotypes, we fitted models including treatment, genotype, and their interaction at each sampling time point. No differences among genotypes were detected at the initial sampling point in November, and there was no effect of temperature or genotype-by-environment interaction at this stage (p > 0.73), indicating similar starting conditions across *six6* genotypes. There was also no evidence for genotype-by-environment interactions on body length, at any sampling point. Although a weak genotype-specific deviation was observed for *six6*\*EL individuals in May (p = 0.049), this effect was small and ended up not being supported by the overall interaction test. There was also no main effect of genotype on body length (p = 0.73), which means that differences in growth among *six6* genotypes are not reflected in this phenotype.

In contrast to body length, we found genotype-dependent differences in growth responses for body weight. While all genotypes showed a strong increase in weight under warm conditions (*p* < 0.001), the magnitude of this response differed among *six6* genotypes, with double homozygotes exhibiting a larger increase compared to heterozygous individuals. Estimated fixed effects showed that heterozygous, *six6*\*EL, individuals gained less weight in the warm treatment relative to six6*EE homozygotes. Overall, individuals with the *six6*\*EE genotype showed a larger increase in body weight between treatments (Δ ≈ 7.2g) compared to *six6*\*EL (Δ ≈ 5.5g) and *six6*\*LL individuals (Δ ≈ 5.8g). Reaction norms differed slightly among genotypes, with *six6*\*EE individuals showing a steeper increase in body weight under warming compared to *six6*\*EL and *six6*\*LL genotypes, indicating a modest gene-by-environment interaction for body weight (Figure 4; *P* = 0.050). No differences among genotypes were observed under control conditions, whereas under warm conditions *six6*\*EE individuals were always significantly heavier than *six6*\*EL individuals (p = 0.006). Further, direct comparison of treatment responses indicated that the increase in body weight was more pronounced for *six6*\*EE than in *six6*\*EL individuals (p = 0.026), while the other genotype contrasts were not significant. A parallel model for body condition in May showed no significant effects of treatment, genotype, or their interaction (p > 0.1). To assess whether the weight G×E was related to overall size, body length was added as a covariate, rendering *six6 × treatment* interaction non-significant (p = 0.36), suggesting that the observed genotype-dependent differences in weight largely reflect variation in overall body size rather than differences in growth independent of length. A parallel model for body condition in May showed no significant effects of treatment, genotype, or their interaction (p > 0.1), indicating consistent condition responses across genotypes. These analyses confirm that fish from the warm treatment were consistently heavier and, most importantly, that the *six6* genotype modified the magnitude of this temperature induced weight increase, with *six6*\*EL genotype showing the smallest gain under warm conditions.

**Figure 4:**
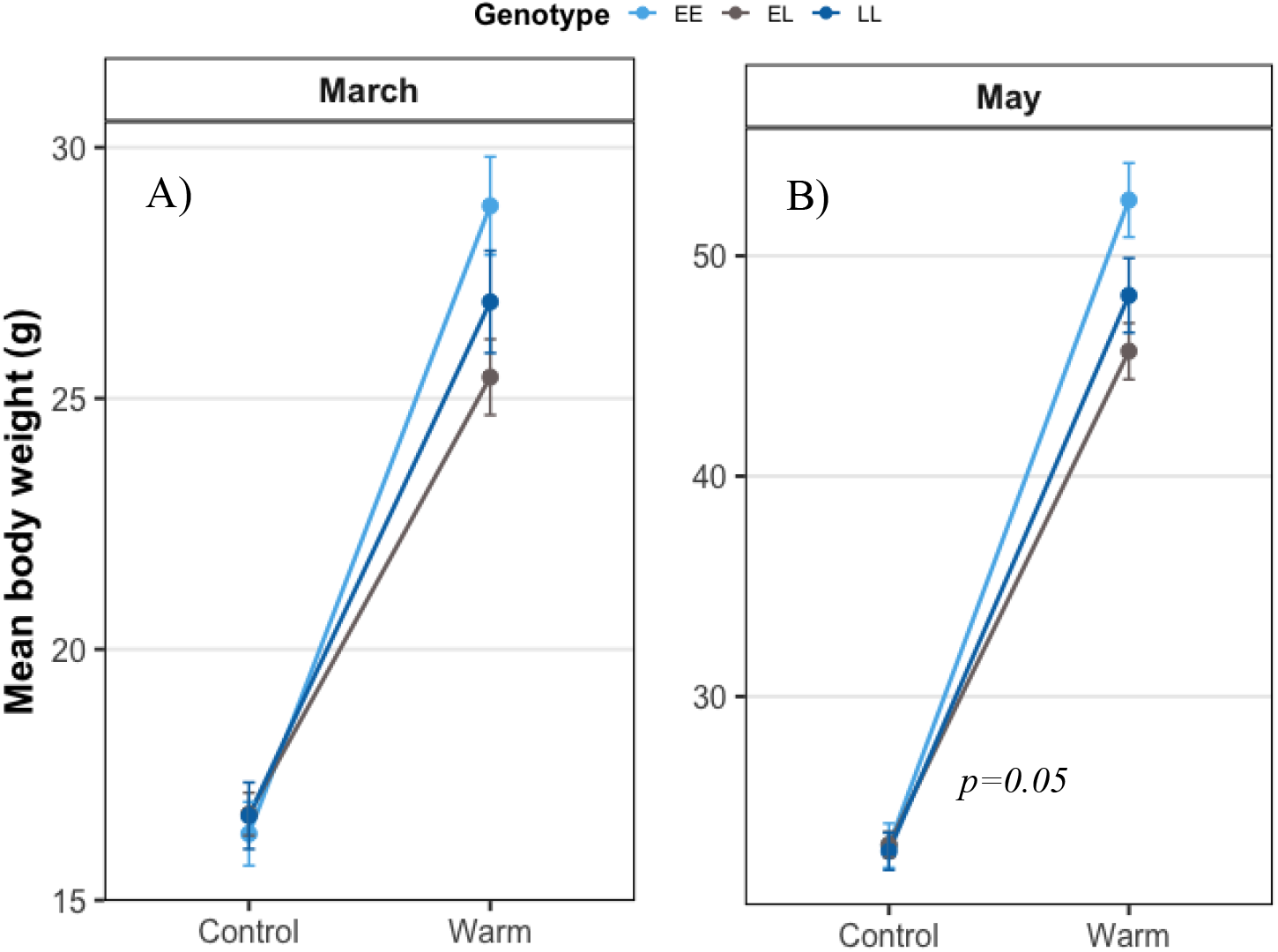
Genotype-specific reaction norms for body weight, under difference thermal regimes in March and May. To explicitly visualize genotype-specific responses to warmer temperature, beyond overall growth patterns, we constructed reaction norms across environments. Mean body weight in March (A) and May (B) of fish reared under control and warm conditions is shown for each *six6* genotype (EE, EL, LL). While all genotypes showed increased body weight under warm conditions, the magnitude of this response differed among genotypes, with *six6*\*EE individuals exhibiting a larger increase compared to *six6\**EL and *six6\**LL individuals. No differences among genotypes were observed under control conditions.

**Figure 5:**
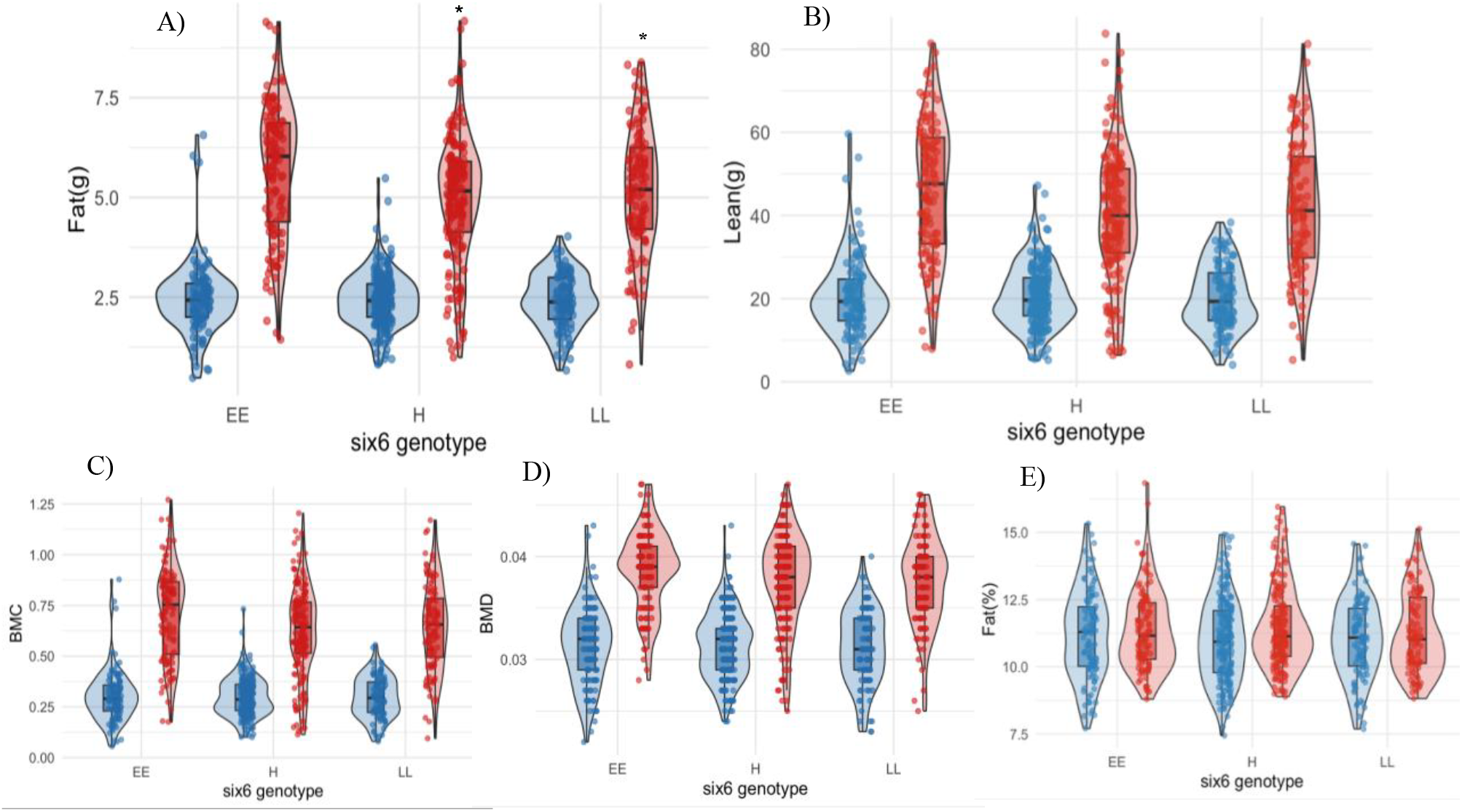
Body composition traits in May across six6 genotypes under control and warm conditions. Violin plots with overlaid boxplots and individual data points are shown for (A) fat mass (g), (B) lean mass (g), (C) bone mineral content (BMC), (D) bone mineral density (BMD), and (E) fat percentage (%). Blue represents control (C) and red represents warm (T) conditions. Across traits, fish reared under warm conditions exhibit higher values overall, with genotype-dependent differences primarily evident in growth-related components (fat mass, lean mass, and BMC), where *six6\**EE individuals show a stronger response to temperature compared to *six6*\*EL and *six6\**LL genotypes. In contrast, proportional traits such as fat percentage show substantial overlap between treatments and genotypes

### 3.3 – Body composition differences dependent on *six6* locus

Body composition traits measured in May, the final sampling point, also showed strong effects of temperature, together with consistent genotype-specific differences for some of the measured traits. Lean (g) and fat (g) mass both showed strong effects of temperature, together with consistent genotype-dependent differences in their response to warmer waters. Lean mass increased substantially under warm conditions (*p* < 0.001), with evidence of a genotype-by-environment interaction (p = 0.052). Heterozygote individuals, *six6*\*EL, showed a reduced response to temperature when compared to *six6*\*EE (*p* = 0.020), with *six6*\*LL individuals showing a similar but still non-significant trend (p = 0.14). Fat mass followed the same general pattern, responding to the temperature treatment (p < 0.001), together with a significant genotype effect (p = 0.0066) and a clear genotype-by-environment interaction (p = 0.0085). Under warm conditions, fat mass significantly increased in the *six6*\*EE individuals (3.19 g), but this increase was reduced in *six6*\*EL (p = 0.0023) and LL individuals (p = 0.037), with a shallower response relative to *six6*\*EE.

Bone-related traits followed a similar pattern, with bone mineral density (BMD) being strongly affected by temperature (p < 0.001) but showing no evidence of genotype effects or G×E (p > 0.36). In contrast, bone mineral content (BMC) exhibited both a significant genotype effect (p = 0.042) and a marginal genotype-by-environment interaction (p = 0.057). Specifically, under warm conditions, *six6*\*EE individuals showed higher BMC relative to six6*EL fish, while *six6*\*LL individuals revealed again the same intermediate values, consistent with patterns observed for both weight and lean mass. Fat percentage was also influenced by the increased temperature environment, with fish in the warm treatment showing higher values of fat, overall (p = 0.0016). There was, however, no evidence for differences among *six6* genotypes (p > 0.3) nor for a genotype-by-environment interaction (p = 0.97), indicating that relative fat allocation was largely similar across genotypes. Taken together, the individual trait analyses point to an important multivariate shift in body composition, which is mostly driven by thermal treatment.

To capture this overall pattern, we performed a principal component analysis (PCA) on the weight-corrected residuals of all the six DXA traits simultaneously (Figure 6).

**Figure 6:**
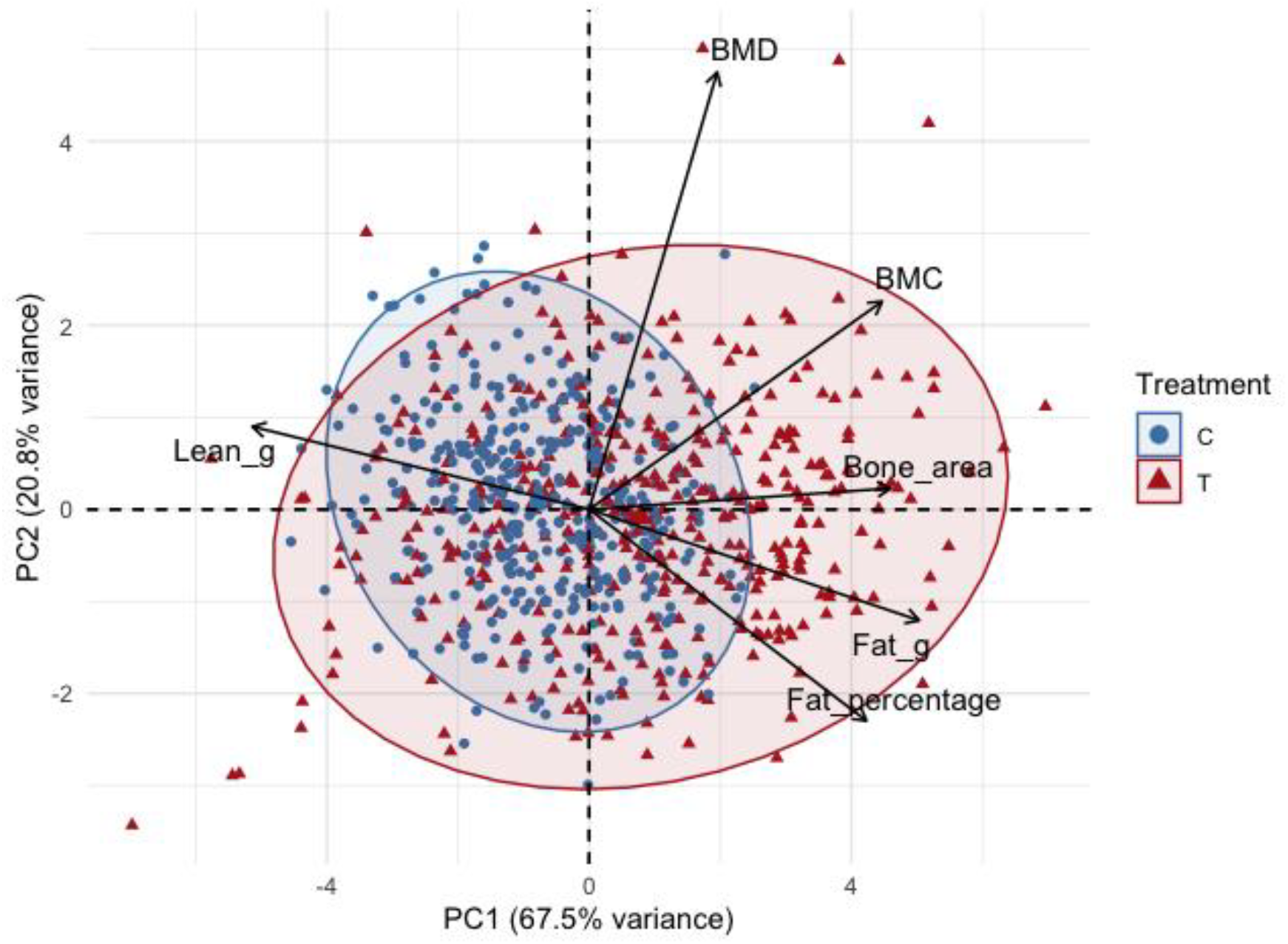
Principal component analysis of weight-corrected body composition traits in control (C) and warm-treated (T) rainbow trout. Each point represents one individual, with blue circles indicating control fish (C) and red triangles indicating warm-treated fish (T). The two principal components shown explain 67.5% (PC1) and 20.8% (PC2) of the total variance in body composition, respectively. PC1 defines a lean-fat trade-off axis, with lean mass (Lean_g) loading negatively and fat mass (Fat_g), fat percentage (Fat_percentage), bone mineral content (BMC) and bone area (Bone_area) loading positively and PC2 captures the independent variation in bone mineral density (BMD). All six DXA-derived traits were residualised against body weight measured in May, to remove size-dependent variation. Arrows indicate the direction and magnitude of each trait’s contribution to the principal components. Shaded ellipses represent 95% confidence regions for each treatment group.

The first two principal components explained 88.3% of the total variance in body composition, with PC1 (67.5%) defining a clear lean vs. fat trade-off axis, with lean mass loading negatively and fat mass, bone area and BMC loading positive, and with PC2 (20.8%) capturing independent variation in bone mineral density. Warm-treated fish were significantly shifted along PC1 relative to controls, reflecting higher relative fat mass and bone size, with lower relative lean mass at similar body weight (R^2^ = 0.115, p = 0.001). Family structure only explained a negligible proportion of the variation (R^2^ < 0.001, p = 0.999), confirming that the observed body composition differences between treatments were not confounded by genetic relatedness among individuals. Together, these results show that the fish developing in warmer waters reallocated their body composition, leading to greater relative fat deposition and bone mineralization, at the expense of lean mass, which was independent of overall body size.

### 3.4. Broad sense heritability of body weight is conserved across environments

Finally, we wanted to understand if the relative importance of kinship, compared to individual-level and tank-level effects, changed under thermal stress. To test it we calculated broad sense heritability (*H*^*2*^) of body weight separately for each thermal environment by partitioning phenotypic variance into family, tank, and residual components using a linear mixed model. Given the lack of an informative pedigree, *H*^*2*^ was calculated as the proportion of phenotypic variance attributable to family identity, hence potentially leading to inflated results, and should be interpreted an estimate capturing both genetic and shared environmental effects. In this design, residual variance serves as a proxy for environmental effects, which due to the absence of a pedigree, cannot be further partitioned. Family identity explained a significant proportion of variation in body weight in both thermal environments (p < 0.001), confirming that body weight has the expected heritable component in this population. Broad-sense heritability was estimated at *H*^*2*^ = 0.29 (95% CI: 0.22–0.36) in control fish and *H*^*2*^ = 0.19 (95% CI: 0.11–0.26) in warm-treated fish. Although the point estimate was lower under warming, the bootstrap confidence interval for the difference between environments included zero (ΔH^2^ = 0.10, 95% CI: 0.00–0.21), we therefore cannot conclude that heritability was significantly reduced by the thermal treatment.

However, a pattern emerged from the variance components themselves, in which the total phenotypic variance (V_total) in body weight was approximately four times greater in the warm environment (317.4) than in controls (82.2). This difference in variance was driven almost entirely by a large expansion of residual variance (V_resid = 255.8 vs 58.5 in controls; Table 1), which we assume as proxy for environmental variation. This suggests that warming substantially increased individual variability in growth, a results that is consistent with greater sensitivity to differences or individual condition under thermal stress. Tank variance was negligible in both environments (Control: 0.00, Warm: 1.23), confirming that tank identity did not confound the treatment effect.

**Table 1.**
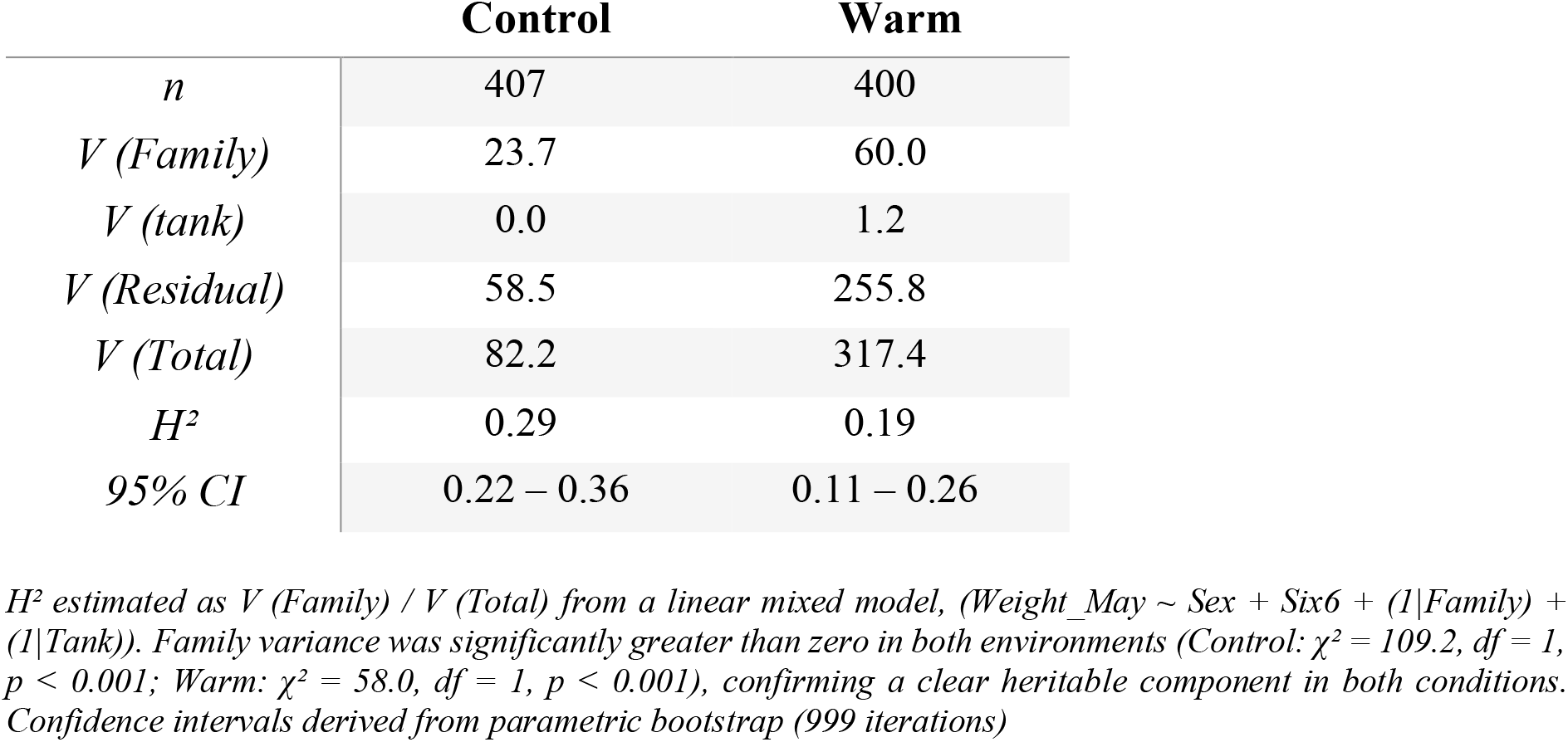
Broad-sense heritability (H^2^) and variance components for body weight, estimated separately for each thermal environment.

## 4. Discussion

As environmental change accelerates, predicting whether and how populations can respond depends critically on understanding how genetic variation at key loci interacts with the environment to shape phenotypic outcomes. The predicted increase in water temperatures is particularly concerning for salmonids given their high oxygen demands and narrow thermal tolerance windows, as warming directly constrains aerobic scope and therefore the capacity to sustain the metabolic costs of growth (Jonsson, 2023). Here, using juvenile rainbow trout reared under contrasting thermal regimes, we show that temperature increase drives a coordinated shift in growth and body composition, and that the magnitude of this response depends in part on genotype at the *six6* locus, a transcription factor repeatedly associated with variation in age at maturity in several salmonid species (Barson et al., 2015; Sinclair-Waters, 2020; 2022; Raunsgard et al., 2023; Waters et al., 2023; Willis et al., 2024). Molecular studies show that *six6* is expressed in the hypothalamus, pituitary gland and retina during development, where it plays a role in regulating the hypothalamic-pituitary-gonadal axis and photoperiod sensing, two pathways closely linked to the coordination of growth and maturation timing in salmonids (Kurko et al., 2020). Consistent with this, our results seem to support the involvement of the *six6* in growth related pathways, under a warming scenario. Given that these pathways are also sensitive to thermal conditions, allelic variation at *six6* may plausibly modulate how individuals partition energy toward growth under thermal stress, making it a strong candidate locus for mediating genotype-dependent responses to warming.

As expected for an ectothermic species, temperature had a strong and consistent effect on growth of the juvenile trout, after being reared for 6 months in a warming scenario (+2°C). Individuals from the warmer environment exhibited substantially higher body weight and length, most likely a reflex from increased metabolic rates and enhanced energy acquisition with more energy allocated to somatic growth (Jonsson 2023). However, and despite the overall significant increase in weight in the warmer temperatures, the responses were not uniform across genotypes. The reduced growth response in *six6*\*EL fish, rather than reflecting the intermediate response typically expected of heterozygotes, highlighted a reduced capacity for plasticity in warming waters when falling below both homozygotes. One possible explanation is related to underdominance, i.e. heterozygote disadvantage, and happens when the expression of two functionally divergent alleles at a large-effect locus produces a less optimal phenotype than each homozygote alone (Soulé, 1979; Livshits and Kobyliansky, 1985). Following this framework, fish that carries both the E and L alleles at the *six6* locus might have distinct, and potentially antagonistic, effects on growth-related pathways which can be partially cancelled or interfere with one another, resulting in a dampened phenotypic response relative to both homozygotes. This interpretation is consistent with the idea that large-effect life-history loci often harbor alleles with different functional consequences, where the heterozygous state might represent a compromise rather than an advantage (Livshits and Kobyliansky, 1985). Importantly, this GxE interaction emerged specifically in the warm treatment, further suggesting that the phenotypic cost of allelic conflict at *six6* is environmentally dependent. One possible explanation lies in the greater metabolic demands imposed by increased temperature, which in turn lead to changes in the regulatory pathways influenced by this locus. Under control conditions, where growth demands are not so strong, the difference between genotypes was not detectable, once again consistent with the idea that genotype-by-environment interactions at functional loci are more likely to be revealed under challenging or novel environmental conditions (Hoffmann and Parsons, 1991).

However, the reduced growth response observed in *six6*\*EL individuals may alternatively reflect a shift in physiological priorities rather than a growth disadvantage per se. Thermal stress in warmer environments often intensifies the competition for aerobic capacity between digestion and other physiological functions in ectotherms. To explain this, Jutfelt et al. (2021) proposed the ‘aerobic scope protection’ hypothesis, describing the voluntary reduction of meal sizes by fish at supra-optimal temperatures to regulate the peak metabolic demand associated with digestion, a process termed specific dynamic action (SDA). At warmer temperatures, the SDA response becomes temporally compressed, generating a higher postprandial peak in oxygen consumption that risks consuming the majority of the animal’s aerobic scope, leaving insufficient residual aerobic capacity for other demanding functions. Under this framework, the slowed growth of *six6*\*EL individuals may not necessarily reflect an inability to grow, but rather a greater reduction in voluntary food intake to protect postprandial residual aerobic scope. This interpretation is further supported by the bioenergetic framework of Huey and Kingsolver (2019), showing that when food intake is reduced, whether through voluntary decrease in food intake or restricted foraging opportunity, both the optimal temperature for growth and the upper thermal limit for growth are lowered. This has consequences that amplify the negative energetic effects of warming by creating what they term as a metabolic meltdown, when accelerating metabolic costs at high temperatures interact with reduced energy intake to dramatically decrease growth. If *six6*\*EL individuals are more prone to voluntary appetite suppression as a protective response under warming, they are experiencing this combined effect, because the energetic cost of feeding under their thermal and aerobic constraints becomes disproportionately high. Under more extreme or prolonged warming, such a conservative strategy could prove advantageous, even at the cost of reduced growth. This is currently supported by theoretical models trying to understand the future impacts of climate change, showing that global warming is often associated with reductions in body size across aquatic systems (Daufresne et al., 2009; Arismendi et al., 2024), potentially driven by physiological constraints such as oxygen limitation (Fenkes et al., 2016).

These same genotype-specific differences were consistently detected across traits associated with overall growth, with the DXA-measured traits showing that *six6* genotype also contributes to variation in body composition, particularly under elevated temperature. While effects on proportional traits such as fat (%) were limited, the absolute measures, like lean mass and BMC revealed consistent genotype differences and a modest G×E, reinforcing the pattern that genotype-dependent growth responses underlie the observed phenotypic variation. Together, our results show that growing up in warmer waters induced an overall change in body composition, shifting fish toward greater relative fat deposition and bone mineralization at the expense of lean tissue, independently of overall body size. Although warm-treated fish were significantly heavier by May, the PCA of weight-corrected residuals revealed that this additional mass was not proportionally distributed across tissue types. The shift in body composition observed in the fish reared in the warmer environment is consistent with temperature dependent responses in other teleosts. When fish are maintained on a diet of fixed macronutrient composition, warmer water increases both feed intake and the rate of lipid digestion and absorption, without a proportional increase in the efficiency of protein retention (Jobling et al, 1997). In another study done with rainbow trout, authors found that rearing the fish at 18°C, compared to 8°C, and feeding them diets of identical protein and energy content, resulted in higher body lipid deposition at the warmer temperature, despite equivalent protein retention, driven by increased feed intake at higher temperatures (Brauge et al, 1995). This suggests that under warming, the additional energy ingested is disproportionately redirected into fat storage rather than lean tissue, even when diet composition remains constant, which is precisely the pattern observed here. This is further supported at the molecular level by studies in other teleosts showing that elevated temperature increases lipid uptake while simultaneously suppressing fat oxidation in muscle tissue (Balbuena-Pecino et al., 2019). In gilthead sea bream, elevated rearing temperatures significantly increased the expression of fatty acid transporters and lipases in muscle while markers of β-oxidation were reduced and key components of the GH/IGF-1 axis were downregulated in both muscle and bone, suggesting impaired musculoskeletal development despite continued overall growth. Together, these mechanisms would produce exactly the compositional pattern we observed with our study, summarized by a pattern of overall growth driven by increased appetite and feed processing, but with disproportionate allocation toward fat rather than lean tissue. The secondary shift toward greater bone mineralization in warm-treated fish, captured along PC2, is consistent with the known relationship between fat mass and bone loading, and may also reflect hormonal co-regulation of adipose and skeletal tissue under altered energy balance, consistent with findings from recent studies on the crosstalk between bone and adipose tissue in zebrafish (Hue et al., 2023). The genotype-dependent differences in lean mass and bone mineral content observed under warming, particularly the attenuated response in heterozygous *six6*\*EL individuals, suggest that the *six6* locus modulates not only overall growth trajectories but also the tissue-specific allocation of growth resources under thermal stress.

Heritable variation in thermal sensitivity of growth has been demonstrated in rainbow trout, with a significant negative genetic correlation between growth rate at lower temperatures and thermal sensitivity across environments, implying that genotypes that perform well in cooler conditions are not necessarily those best suited to warmer ones (Janhunen et al., 2016). The presence of a clear GxE in body weight, together with differences in body composition traits, prompted us to ask whether these responses have a detectable genetic basis in our population. Lacking a fully resolved pedigree, which would allow the calculation of proper estimates of additive genetic and environmental variance components, i.e. narrow-sense heritability, we asked whether broad-sense heritability could provide ‘reliable intuition’ on the importance of genetic causes in producing phenotypic variation an approach that, while limited, can offer meaningful insight into the relative contribution of family-level resemblance to trait variation (Tal, 2012). Even when using a limited calculation, we still confirmed the contribution of family in explaining a significant, and substantial, proportion of body weight variation in both thermal environments. Our broad-sense heritability estimates are consistent with narrow-sense estimates reported for growth traits in rainbow trout reared at elevated temperatures in other studies, with values of 0.24 for to thermal sensitivity of growth (Janhunen et al., 2016) and 0.41 for growth under chronic thermal stress (Gallardo-Hidalgo et al., 2021). The consistency between our broad-sense H^2^ estimates and narrow-sense h^2^ values from fully resolved pedigree studies suggests that our family-based approach cwas capable of capturing a meaningful signal of true additive genetic variance, despite the absence of a full pedigree.Although the point estimate was lower under warming, the confidence interval for the difference between environments included zero, and we therefore cannot conclude that heritability itself was significantly reduced by the thermal treatment. In the absence of a fully resolved pedigree, these estimates capture additive genetic variance alongside dominance, maternal effects, and shared early environmental effects, and should be interpreted as upper-bound broad-sense values rather than narrow-sense heritability (Lynch and Walsh, 1998).

However, the most striking pattern in the variance decomposition was not the H^2^ result itself, but the approximately four-fold expansion of total phenotypic variance under warming, almost entirely driven by residual variance, which in our design serves as a proxy for environmental sensitivity. Early studies of heritability under contrasting environmental conditions showed that the magnitude of genetic and environmental components of phenotypic variation can change with environmental quality, with unfavorable or novel conditions often increasing environmental variance relative to genetic variance, thereby reducing heritability even when absolute genetic variance remains stable (Falconer, 1952; Hoffmann and Merilä, 1999). The pattern observed here is consistent with this framework, we see that warming did not eliminate the heritable component of body weight but substantially amplified the individual-level variance around it, suggesting that families, or individuals of each family, responded differently to the thermal challenge depending on their individual condition. This is well supported by the knowledge that organism in stress-inducing environments show higher phenotypic variance, and it has been interpreted as reflecting the reduced developmental integration under challenging conditions. This gives rise to more individual differences resulting from stress responsiveness, that become phenotypically expressed in ways that are buffered in benign environments (Hoffmann and Parsons, 1991). Whether this expanded individual variance has a heritable component. i.e. some families are consistently more variable than others under warming, cannot be resolved with the current design but represents an important question for future work using a full pedigree and animal model approach, as well as a bigger sample size in both number of families and individuals.

It is important to note that our experimental conditions represent an environment with abundant food availability, with fish being fed *ad libitum*, which in unlikely to occur in the wild. In natural systems, the increase in water temperature will most likely not coincide with increased resource availability (Daufresne et al., 2009; Jeppesen et al., 2012) and can even exaggerate any underlying energetic constraints. Under such conditions, genotypes with lower growth rates but reduced energetic demands could potentially have a relative advantage. Also, our experiment covered only a six-month window of early life, from six months to one year of age, so remains unclear if the growth and body composition differences observed here translate into fitness consequences over longer timescales. Short-term common-garden experiments are well suited to detecting plastic responses and genotype-dependent reaction norms, but determining whether such plasticity is adaptive, maladaptive, or neutral ultimately requires linking phenotypic responses to survival and reproductive success across full life histories. In salmonids, this is particularly challenging given their complex life histories, where conditions experienced during early freshwater residence can have carry-over effects on marine performance and reproductive timing that are difficult to predict from juvenile phenotypes alone (Crozier et al. 2008). Furthermore, phenotypic mismatches, where plastic responses shift phenotypes in a direction that is initially beneficial but becomes maladaptive as environmental conditions continue to change, represent a real risk under scenarios of sustained warming and are expected to lead to local extinction (Urban, 2015). Whether the genotype-dependent growth and shifts resulting from our study reveal a form of adaptive response to a of a novel thermal environment or leads to energetic reallocation that may compromise downstream fitness, remains to be further investigated and is an important avenue for future work. Regardless, our findings contribute to a growing body of evidence that the effects of large-effect life-history loci are highly dependent on the environment, by mediating how individuals translate environmental signals into phenotypic outcomes, with potentially important consequences for population-level adaptation under climate warming.

## References

Arismendi, I., Gregory, S.V., Bateman, D.S., and Penaluna, B.E. (2024). Shrinking sizes of trout and salamanders are unexplained by climate warming alone. Scientific Reports, 14, 13614. 10.1038/s41598-024-64145-x

Aykanat, T., McLennan, D., Metcalfe, N.B., and Prokkola, J.M. (2024). Early survival in Atlantic salmon is associated with parental genotypes at loci linked to timing of maturation. Evolution, 78(8), 1441–1452. 10.1093/evolut/qpae072

Åsheim, E.R.; Debes, P.V.; House, A.; Liljeström, P.; Niemelä, P.T.; Siren, J.P.; Erkinaro, J. and Primmer, C.R. (2021) Atlantic salmon (Salmo salar) age at maturity is strongly affected by temperature, population and age-at-maturity genotype. Conservation Physiology, 11, 10.1093/conphys/coac086

Balbuena-Pecino, S., Riera-Heredia, N., Vélez, E.J., Gutiérrez, J., Navarro, I., Riera-Codina, M., & Capilla, E. (2019). Temperature affects musculoskeletal development and muscle lipid metabolism of gilthead sea bream (Sparus aurata). Frontiers in Endocrinology, 10, 173. 10.3389/fendo.2019.00173

Barson, N.J., Aykanat, T., Hindar, K., Baranski, M., Bolstad, G.H., Fiske, P., Jacq, C., Jensen, A.J., Johnston, S.E., Karlsson, S., Kent, M., Moen, T., Niemelä, E., Nome, T., Næsje, T.F., Orell, P., Romakkaniemi, A., Sægrov, H., Urdal, K., Erkinaro, J., Lien, S., and Primmer, C.R. (2015). Sex-dependent dominance at a single locus maintains variation in age at maturity in salmon. Nature, 528, 405–408. 10.1038/nature16062

Brauge, C., Corraze, G., & Médale, F. (1995). Effect of dietary levels of lipid and carbohydrate on growth performance, body composition, nitrogen excretion and plasma glucose levels in rainbow trout reared at 8 or 18°C. Reproduction, Nutrition, Development, 35(3), 277–290. 10.1051/rnd:19950304

Cauwelier, E., Gilbey, J., Sampayo, J., Stradmeyer, L. and Middlemas, S.J. (2018). Identification of a single genomic region associated with seasonal river return timing in adult Scottish Atlantic salmon (Salmo salar), using a genome-wide association study. Canadian Journal of Fisheries and Aquatic Sciences, 75(9), 1427–1435. 10.1139/cjfas-2017-0293

Chevin, L.M., Lande, R. and Mace, G.M. (2010). Adaptation, plasticity, and extinction in a changing environment: towards a predictive theory. PLoS Biology, 8(4), e1000357. 10.1371/journal.pbio.1000357

Crozier, L.G. and Hutchings, J.A. (2014). Plastic and evolutionary responses to climate change in fish. Evolutionary Applications, 7(1), 68–87. 10.1111/eva.12135

Daufresne, M., Lengfellner, K. and Sommer, U. (2009). Global warming benefits the small in aquatic ecosystems. Proceedings of the National Academy of Sciences, 106(31), 12788– 12793.10.1073/pnas.0902080106

Doctor, K., Berejikian, B., Hard, J. J., and VanDoornik, D. (2014). Growth-Mediated life history traits of Steelhead reveal phenotypic divergence and plastic response to temperature. Transactions of the American Fisheries Society, 143(2), 317–333. 10.1080/00028487.2013.849617

Dubey, M.K., Kamalam, B.S., Rajesh, M., Sarma, D., Pandey, A., Baral, P., and Sharma, P. (2023). Exposure to different temperature regimes at early life stages affects hatching, developmental morphology, larval growth, and muscle cellularity in rainbow trout, Oncorhynchus mykiss. Fish Physiology and Biochemistry, 49(2), 219–238. 10.1007/s10695-023-01166-9

Falconer, D.S. and Mackay, T.F.C. (1996). Introduction to Quantitative Genetics (4th ed.). Longman.

Fenkes, M., Shiels, H.A., Fitzpatrick, J.L. and Nudds, R.L. (2016). The potential impacts of migratory difficulty, including warmer waters and altered flow conditions, on the reproductive success of salmonid fishes. Comparative Biochemistry and Physiology Part A: Molecular & Integrative Physiology, 193, 11–21. 10.1016/j.cbpa.2015.11.012

Friedland, K.D., Chaput, G., & MacLean, J.C. (2005). The emerging role of climate in post-smolt growth of Atlantic salmon. ICES Journal of Marine Science, 62(7), 1338–1349. 10.1016/j.icesjms.2005.04.012

Gallardo-Hidalgo, J., Barría, A., Yoshida, G.M. and Yáñez, J.M. (2021). Genetics of growth and survival under chronic heat stress and trade-offs with growth- and robustness-related traits in rainbow trout. Aquaculture, 531, 735685.10.1016/j.aquaculture.2020.735685

Hoffmann, A.A., & Sgrò, C.M. (2011). Climate change and evolutionary adaptation. Nature, 470, 479–485. 10.1038/nature09670

Hoffmann, A.A. and Merilä, J. (1999). Heritable variation and evolution under favourable and unfavourable conditions. Trends in Ecology and Evolution, 14(3), 96–101. 10.1016/S0169-5347(99)01595-7

Hoffmann, A.A., and Parsons, P.A. (1991). Evolutionary Genetics and Environmental Stress. Oxford University Press.

Huey, R.B., and Kingsolver, J.G. (2019). Climate warming, resource availability, and the metabolic meltdown of ectotherms. American Naturalist, 194, E140–E150. 10.1086/705679

IPCC. 2024. Climate Change 2024: Impacts, Adaptation and Vulnerability. Cambridge University

Janhunen, M., Kause, A., Vehviläinen, H. and Järvisalo, O. (2016). Thermal sensitivity of growth indicates heritable variation in 1-year-old rainbow trout (Oncorhynchus mykiss). Genetics Selection Evolution, 48, 77. 10.1186/s12711-016-0272-3

Jobling, M. (1997). Temperature and growth: modulation of growth rate via temperature change. In C.M. Wood & D.G. McDonald (Eds.), Global Warming: Implications for Freshwater and Marine Fish (pp. 225–253). Cambridge University Press.

Jonsson, B. and Jonsson, N. (2009). A review of the likely effects of climate change on anadromous Atlantic salmon Salmo salar and brown trout Salmo trutta, with particular reference to water temperature and flow. Journal of Fish Biology, 75(10), 2381–2447. 10.1111/j.1095-8649.2009.02380.x

Jonsson, B. and Jonsson, N. (2014). Early environment influences later performance in fishes. Journal of Fish Biology, 85(2), 151–188. 10.1111/jfb.12432

Jonsson, B. (2023). Thermal effects on ecological traits of salmonids. Fishes, 8(7), 337. 10.3390/fishes8070337

Jutfelt, F., Norin, T., Åsheim, E.R., Rowsey, L.E., Andreassen, A.H., Morgan, R., Clark, T.D., and Speers-Roesch, B. (2021). ‘Aerobic scope protection’ reduces ectotherm growth under warming. Functional Ecology, 35, 1397–1407. 10.1111/1365-2435.13811

Kurko, J., Debes, P.V., House, A.H., Aykanat, T., Erkinaro, J. and Primmer, C.R. (2020). Transcription profiles of age-at-maturity-associated genes suggest cell fate commitment regulation as a key factor in the Atlantic salmon maturation process. G3: Genes, Genomes, Genetics, 10(1), 235–246. 10.1534/g3.119.400882

Livshits, G. and Kobyliansky, E. (1985). Lerner’s concept of developmental homeostasis and the problem of heterozygosity level in natural populations. Heredity, 55, 341–353. 10.1038/hdy.1985.117

Le Coeur, C., Yoccoz, N.G., Salguero-Gómez, R. and Vindenes, Y. (2022). Life history adaptations to fluctuating environments: Combined effects of demographic buffering and lability. Ecology Letters, 25(10), 2107–2119. 10.1111/ele.14071

Lynch, M. and Walsh, B. (1998). Genetics and Analysis of Quantitative Traits. Sinauer Associates.

Mackay, T.F.C., Stone, E.A., & Ayroles, J.F. (2009). The genetics of quantitative traits: challenges and prospects. Nature Reviews Genetics, 10(8), 565–577. 10.1038/nrg2612

Merilä, J. and Hendry, A.P. (2014). Climate change, adaptation, and phenotypic plasticity: the problem and the evidence. Evolutionary Applications, 7(1), 1–14. 10.1111/eva.12137

Moustakas-Verho, J.E., Kurko, J., House, A.H., Erkinaro, J., Debes, P. and Primmer, C.R. (2020). Developmental expression patterns of six6: A gene linked with spawning ecotypes in Atlantic salmon. Gene Expression Patterns, 38, 119149. 10.1016/j.gep.2020.119149

Oomen, R.A. and Hutchings, J.A. (2022). Genomic reaction norms inform predictions of plastic and adaptive responses to climate change. Journal of Animal Ecology, 91(6), 1073– 1087. 10.1111/1365-2656.13707

Pashay Ahi, E, Lindeza A S, Miettinen, A and Primmer, C (2025) ‘Transcriptional responses to changing environments: insights from salmonids’, Reviews in Fish Biology and Fisheries, vol. 35, no. 2, pp. 681–706. doi.org/ 10.1007/s11160-025-09928-9

Parmesan, C. (2006). Ecological and evolutionary responses to recent climate change. Annual Review of Ecology, Evolution, and Systematics, 37, 637–669. 10.1146/annurev.ecolsys.37.091305.110100

Pörtner, H.O. and Farrell, A.P. (2008). Physiology and climate change. Science, 322(5902), 690–692. 10.1126/science.1163156

Raunsgard, A., Persson, L., Czorlich, Y., Ugedal, O., Thorstad, E. B., Karlsson, S., Fiske, P. and Bolstad, G. H. (2024). Variation in phenotypic plasticity across age-at-maturity genotypes in wild Atlantic salmon. Molecular Ecology, 33(3), e17229. 10.1111/mec.17229

Reed, T.E., Schindler, D.E. and Waples, R.S. (2010). Interacting effects of phenotypic plasticity and evolution on population persistence in a changing climate. Conservation Biology, 24(1), 317–325. 10.1111/j.1523-1739.2009.01378.x

Russell, I.C., Aprahamian, M.W., Barry, J., Davidson, I.C., Fiske, P., Ibbotson, A.T., Kennedy, R.J., Maclean, J.C., Moore, A., and Otero, J. (2012). The influence of the freshwater environment and the biological characteristics of Atlantic salmon smolts on their subsequent marine survival. ICES Journal of Marine Science, 69(9), 1563–1573. 10.1093/icesjms/fsr208

Sae-Lim, P., Kause, A., Mulder, H.A., Martin, K.E., Barfoot, A.J., Parsons, J.E., Davidson, J., Rexroad, C.E., van Arendonk, J.A.M., & Komen, H. (2013). Genotype-by-environment interaction of growth traits in rainbow trout (Oncorhynchus mykiss): a continental scale study. Journal of Animal Science, 91(12), 5572–5581. 10.2527/jas.2012-5949

Sae-Lim, P., Kause, A., Janhunen, M., Vehviläinen, H., Koskinen, H., Gjerde, B., Lillehammer, M., and Mulder, H.A. (2015). Genetic (co)variance of rainbow trout (Oncorhynchus mykiss) body weight and its uniformity across production environments. Genetics Selection Evolution, 47, 46. 10.1186/s12711-015-0122-8

Siitonen, L. (1986). Factors affecting growth in rainbow trout (Salmo gairdneri) stocks. Aquaculture, 57(1–4), 185–191. 10.1016/0044-8486(86)90196-1

Sinclair-Waters, M., Nome, T., Wang, J., Lien, S., Kent, M.P., Sægrov, H., Florø-Larsen, B., Bolstad, G.H., Primmer, C.R., and Barson, N.J. (2022). Dissecting the loci underlying maturation timing in Atlantic salmon using haplotype and multi-SNP based association methods. Heredity, 129(6), 356–365. 10.1038/s41437-022-00570-w

Sinclair-Waters, M., Ødegård, J., Korsvoll, S. A., Moen, T., Lien, S., Primmer, C. R. and Barson, N. J. (2020). Beyond large-effect loci: Large-scale GWAS reveals a mixed large-effect and polygenic architecture for age at maturity of Atlantic salmon. Genetics Selection Evolution, 52(1), 9. 10.1186/s12711-020-0529-8

Soulé, M.E. (1979). Heterozygosity and developmental stability: another look. Evolution, 33(1), 396–401. 10.1111/j.1558-5646.1979.tb04693.x

Tal, O. (2012). The impact of gene–environment interaction and correlation on the interpretation of heritability. Acta Biotheoretica, 60(3), 225–237. 10.1007/s10441-011-9139-8

Tuljapurkar, S., Steiner, U.K and Orzack, S. H. (2009). Dynamic heterogeneity in life histories. Ecollogy Letters;12:93–106. doi: 10.1111/j.1461-0248.2008.01262.x

Urban, M.C. (2015). Accelerating extinction risk from climate change. Science, 348(6234), 571–573. 10.1126/science.aaa4984

Vehanen, T., Sutela, T., and Huusko, A. (2023). Potential impact of climate change on salmonid smolt ecology. Fishes, 8(7), 382. 10.3390/fishes8070382

Volkoff, H., & Rønnestad, I. (2020). Effects of temperature on feeding and digestive processes in fish. Temperature (Austin), 7(4), 307–320. 10.1080/23328940.2020.1765950

Waters, C. D., Clemento, A., Aykanat, T., Garza, J. C., Naish, K. A., Narum, S., and Primmer, C. R. (2021). Heterogeneous genetic basis of age at maturity in salmonid fishes. Molecular Ecology, 30(6), 1435–1456.10.1111/mec.15822

Willis, S., Stephenson, J., Pierce, A., Medeiros, L., Jenkins, L., Hatch, D. R., and Narum, S. (2024). A genomic region associated with iteroparous spawning phenology is linked with age-at-maturity in female steelhead trout. Evolutionary Applications, 17(2), e13622. 10.1111/eva.13622

Wilkes, D., Xie, S.Q., Stickland, N.C., Alami-Durante, H., Kentouri, M., Sterioti, A., Koumoundouros, G., Fauconneau, B. and Goldspink, G. (2001). Temperature and myogenic factor transcript levels during early development determines muscle growth potential in rainbow trout (Oncorhynchus mykiss) and sea bass (Dicentrarchus labrax). Journal of Experimental Biology, 204(16), 2763–2771. 10.1242/jeb.204.16.2763

